# Protein solubility depends on centrifugation: Aiki-Sol, a per-regime predictor for *E. coli*

**DOI:** 10.64898/2026.05.14.725067

**Authors:** Riya Rajagopalan, Radheesh Sharma Meda, Shankar Shastry, Venkatesh Mysore

## Abstract

**Motivation:** Sequence-based predictors of recombinant protein solubility in *Escherichia coli* have plateaued (NESG independent-test AUC 0.760 →~ 0.80 over eight years of protein-language-model variants). The plateau hides a latent confound: the centrifugation regime used to separate the soluble from the insoluble fraction is a hidden variable collapsed into a single binary “soluble” label. The protein’s biochemistry does not change between regimes; what changes is which fraction of the lysate is recovered as soluble. Existing predictors treat the regime as label noise rather than a feature, and sequence overlap between training and test partitions masks the resulting failure mode.

**Results:** We release the Aiki-Sol Dataset, a tiered *E. coli* solubility corpus: a ~ 85K stringency-annotated benchmark, an Apache-licensed ~ 147K extension adding binary-only-labelled proteins, and a ~ 229K research-tier pool incorporating non-commercially-licensed sources. On the ~ 85K benchmark, scored on sequence-cluster-disjoint partitions, the strongest published binary comparator falls below chance on the 32,000 × *g* stratum (AUC 0.491 ± 0.020); a fine-tuned ESM-2 650M backbone with five protocol-matched outputs lifts pooled AUC by +0.108 (paired-bootstrap CI lower bound ≥ +0.090). The gain is curation, not architecture: structure-aware predictors given ESMFold structures do not outperform the sequence-only frame, and capacity scaled to 3B parameters does not exceed the conditioned 650M backbone. The released model, Aiki-Sol, jointly supervises five per-stringency outputs alongside a marginal output for stringency-unknown proteins; on five external cohorts it lifts cohort-mean AUC from 0.69–0.70 to 0.825, with a ≥ +0.10–0.16 lift on the three cohorts at measurably-zero training-pool overlap.

**Availability and implementation:** Aiki-Sol model weights (Apache 2.0), the 147K-row license-clean training pool of the deployment checkpoint (CC BY 4.0), the cluster-disjoint per-stringency 5-fold partition assignments, per-cohort prediction CSVs, and source code for training, inference, and figure reproduction are available at https://github.com/aikium-public/aiki-sol and archived at Zenodo 10.5281/zenodo.20151817. The research-tier 229K checkpoint is released under CC-BY-NC-ND 4.0 (inheriting the most-restrictive upstream-source tier); its training CSV and the 84,809-protein stringency-annotated benchmark of §2.1 mix non-commercial-tier upstream sources and are not redistributed verbatim. Upstream sources are documented in Data availability and SI §S1. The deployment artefact is distributed as a Python package (`pip install aikisol`) with a `predict(seq)` entry point.

**Supplementary information:** Supplementary text, figures, and tables are available at *Bioinformatics* online.

## 1 Introduction

Insolubility of proteins during recombinant expression in *Escherichia coli* or in subsequent purification steps wastes weeks of experimental effort; computational solubility prediction before synthesis can improve hit rates [1] and is the standard pre-commitment filter in industrial protein-engineering pipelines. Sequence-based predictors have evolved through eight years of protein-language-model variants [3, 8, 11, 12, 13, 17, 21, 22], yet reported NESG independent-test AUC has moved only from 0.760 [3] to roughly 0.80 [19, 20].

Two structural problems with this literature are under-acknowledged. The first is a latent confound in the training data: the centrifugation regime used to separate the soluble from the insoluble fraction is a hidden variable that has been collapsed into a single binary “soluble” column. The protein’s biophysical state does not change between regimes; what changes is which fraction of the lysate is recovered as soluble. A 3,000 × *g* for 10 min spin is a debris-clearing clarification [6, 5]; 32,000 × *g* additionally clears aggregated species; 100,000 × *g* pellets ribosomes and large soluble multimers in addition to aggregates [7]; and the cell-free eSol/PURE system at 21,600 × *g* [4] is likely correlated with chaperone-independent intrinsic aggregation propensity [23]. Per-protocol soluble fractions across these assays span 65%–99.7% on overlapping protein populations (Fig. 1); collapsing the resulting labels into a single column, as existing predictors do, conflates five operationally distinct phenotypes that a protein scientist needs to keep separate, because different purification, assay, and formulation workflows are bottlenecked by different regimes.

**Figure 1:**
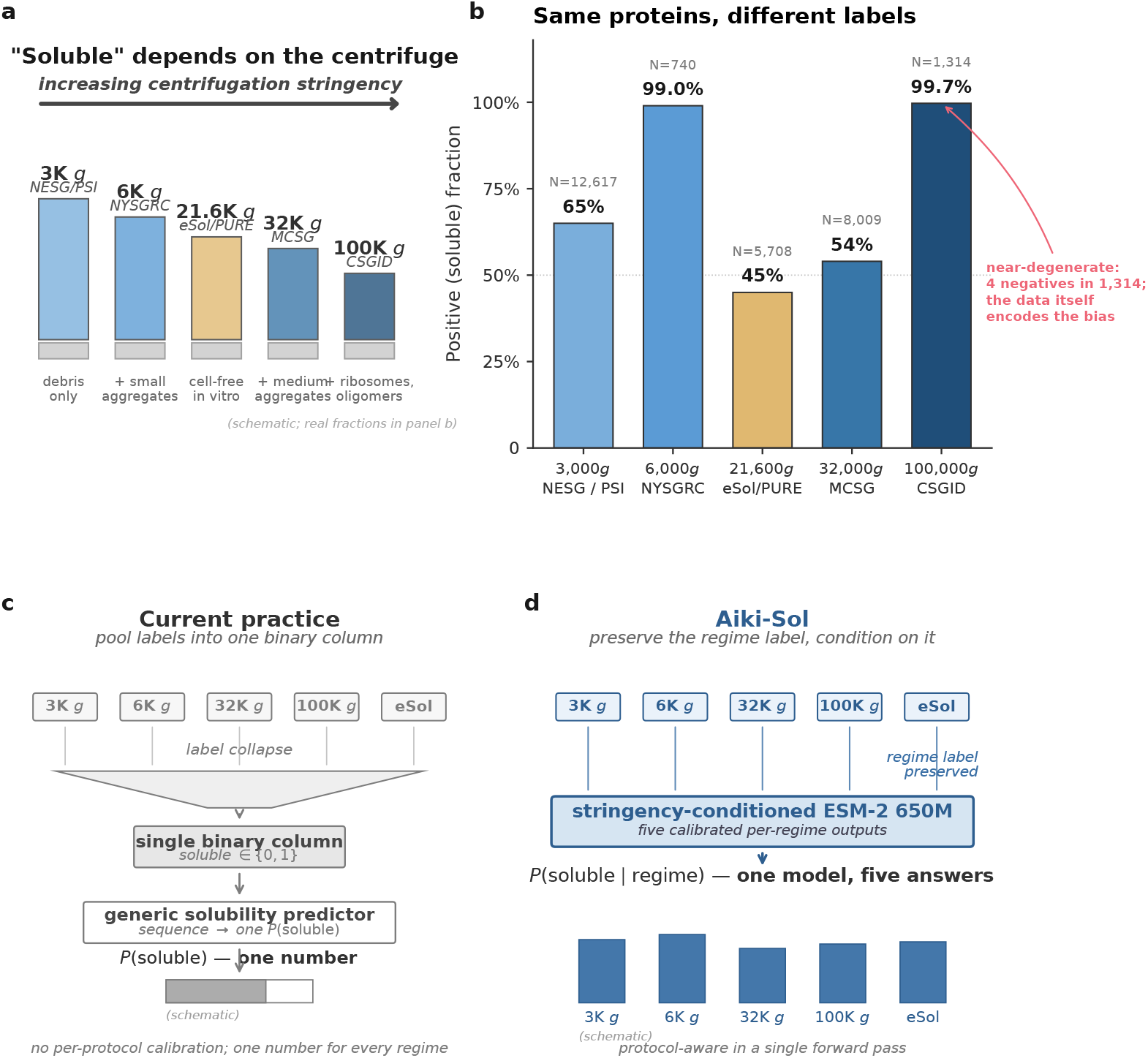
“Soluble” is not one phenotype, and what to do about it. **(a)** Each public consortium operationalises “soluble” at a different centrifugation g-force, clearing a different subpopulation of the lysate: NESG’s 3,000*g ×* 10 min spin clears cellular debris; 32,000*g* additionally clears aggregated species; 100,000*g* pellets ribosomes and large multimers; the cell-free eSol/PURE system at 21,600*g* reports chaperone-independent intrinsic aggregation propensity in vitro. Bar heights are schematic. **(b)** On the protocol-stratified pool, the same protein populations carry positive (soluble) fractions spanning 45%–99.7%, set only by which centrifugation regime was used. Per-center g-force mappings (HIGH-confidence: NESG, NYSGRC, CESG; MEDIUM, rotor inferred: MCSG, CSGID, EFI) are documented in Methods §S1. **(c)** Current practice pools the five regimes into one binary column and trains a generic predictor that emits one *P* (soluble) with no record of which regime it refers to. **(d)** Aiki-Sol preserves the regime as a per-protein label and conditions a single fine-tuned ESM-2 650M checkpoint on it, emitting five calibrated per-regime probabilities {*P* (soluble | *g*)}_*g*_ in one forward pass. The five per-regime rankings agree closely (mean pairwise Spearman *ρ* = 0.972, §2.2); the model’s job is to set where the soluble/insoluble cut falls for each regime, a deployment property no published predictor offers.

The second problem is sequence-level overlap between training and test partitions. A test protein sharing ≥ 25% pairwise identity with a training-pool sequence cannot establish generalisation to genuinely unseen sequence space; cluster-disjoint evaluation, standard for protein-function benchmarks [10], has not been routinely applied to cross-predictor comparison of *E. coli* solubility models. The two problems compound: a binary predictor trained on a pool that under-represents one centrifugation regime will fail on that regime at deployment, and the failure stays hidden under evaluation against partly-overlapping test sets, where resemblance to a training sequence can substitute for a genuine prediction.

We address both problems jointly: a tiered *E. coli* solubility corpus (Table 2) and a single fine-tuned ESM-2 650M backbone that emits five protocol-matched probabilities per protein in one forward pass. We benchmark against seven published predictors [3, 12, 11, 8, 17, 13, 14] on cluster-disjoint partitions and release data, partitions, and code under permissive licenses.

## 2 Results

### 2.1 Cluster-disjoint per-stringency 5-fold benchmark

We benchmark Aiki-Sol against the strongest published binary predictors on the protocol-stratified five-fold cluster-disjoint partition of the 84,809-protein protocol-stratified pool (§4.3; sequence clustering at 25% identity, 80% coverage, ~ 17,000 test proteins per fold spanning all five centrifugation regimes). Seven published predictors are scored on the same test proteins using the authors’ released checkpoints and inference code (no retraining or removal of their training data from our test folds): `PLM_Sol` [12], `NetSolP` [3], SoluProt [11], DeepSol1 [8], DSResSol [17], and the two structure-aware predictors GATSol [13] and ProtSolM [14].

The four Aiki-Sol variants in Table 1 share the fine-tuned ESM-2 650M backbone and differ only in head/loss specification (per-head specs in §4.6, SI S4). The benchmark uses the protocol-stratified 84,809-protein pool of Table 2, on which all seven comparators have prior cluster-disjoint 5-fold scoring. The deployment checkpoint’s parallel cluster-disjoint 5-fold benchmark on canonical-147K (comparator predictions re-binned under the canonical fold assignment) appears in Table S6, and cross-pool generalisation on five external cohorts in Table S5.

**Table 1:**
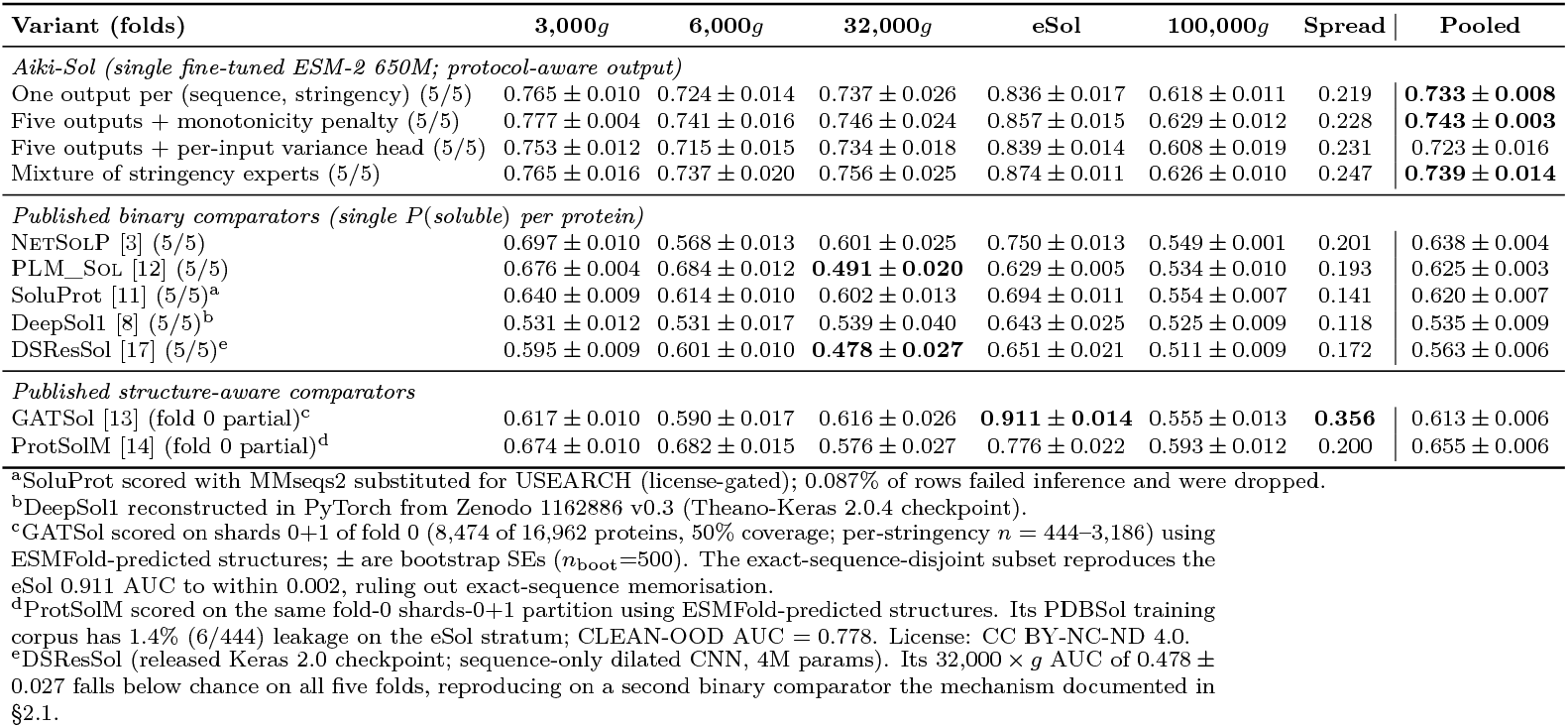
Cluster-disjoint per-stringency AUC on the protocol-stratified five-fold test partition (~17,000 proteins per fold, 84,809 total across folds; sequence clustering at 25% identity, 80% coverage). Aiki-Sol variants emit five protocol-matched probabilities per protein (one forward pass); for binary comparators the same predicted probability is scored against each protein’s stringency-matched binary label. Mean ± sample standard deviation across folds; the “Spread” column is the per-variant max−min stringency AUC.

**Table 2:** Pool taxonomy. Row counts re-verified on disk. “Used by” references the manuscript section / table where the pool’s numbers appear. The two trained pools (canonical-147K, research-tier 229K) supply the released deployment checkpoint and the research-tier higher-coverage checkpoint respectively; the protocol-stratified 84.8K pool supports the headline cluster-disjoint per-stringency 5-fold benchmark of §2.1.

| Pool name | Construction | Rows | License tier | Used by |
| --- | --- | --- | --- | --- |
| lineage core | strict-curated *E. coli* subset | 35,808 | mixed | capacity probe (§2.2); early training splits |
| protocol-stratified 84.8K | known-stringency, permissively-licensed subset of the curated literature collection (mixes Apache + CC-BY-NC-ND-tagged sources) | 84,809 | mixed | §2.1 headline 5-fold benchmark |
| canonical-147K | Apache-tier curated union | 147,574 | Apache 2.0 | Aiki-Sol training (§4.6) |
| research-tier 229K | canonical-147K + CC-BY-NC-ND sources | 229,349 | CC-BY-NC-ND 4.0 | Aiki-Sol-research checkpoint |

#### Surfacing the centrifugation-regime confound lifts AUC by +0.095 to +0.208 over the published binary comparators

(Figure 2; Table 1). The single design choice that produces this lift is straightforward: the model is told which centrifugation regime each prediction is for, so the regime stops hiding inside the binary label. Every Aiki-Sol variant pools above 0.733; the strongest published comparators reach `NetSolP` 0.638 ± 0.004, `PLM_Sol` 0.625 ± 0.003, ProtSolM 0.655 ± 0.006, and the remaining four (SoluProt, DeepSol1, DSResSol, GATSol) sit between 0.535 and 0.620. The +0.108 gap of the conditioned baseline over `PLM_Sol` is an order of magnitude larger than the spread among Aiki-Sol variants (±0.005 to ±0.015), and a 2,000-iteration paired bootstrap confirms a positive lift over every binary comparator individually (narrowest ΔAUC CI_95_ lower bound +0.090, vs. `PLM_Sol`); what matters is that the model is told the regime, not the specific output-head architecture on top. On the protocol-stratified cluster-disjoint partitions used here, no published binary comparator from 2018–2025 pools above 0.66. The +0.108 gap is conservative: any unmeasured leakage between a comparator’s training pool and our test folds would inflate the comparator’s AUC. External cohorts with Aiki-Sol’s training-pool overlap measured is discussed in §2.3.

**Figure 2:**
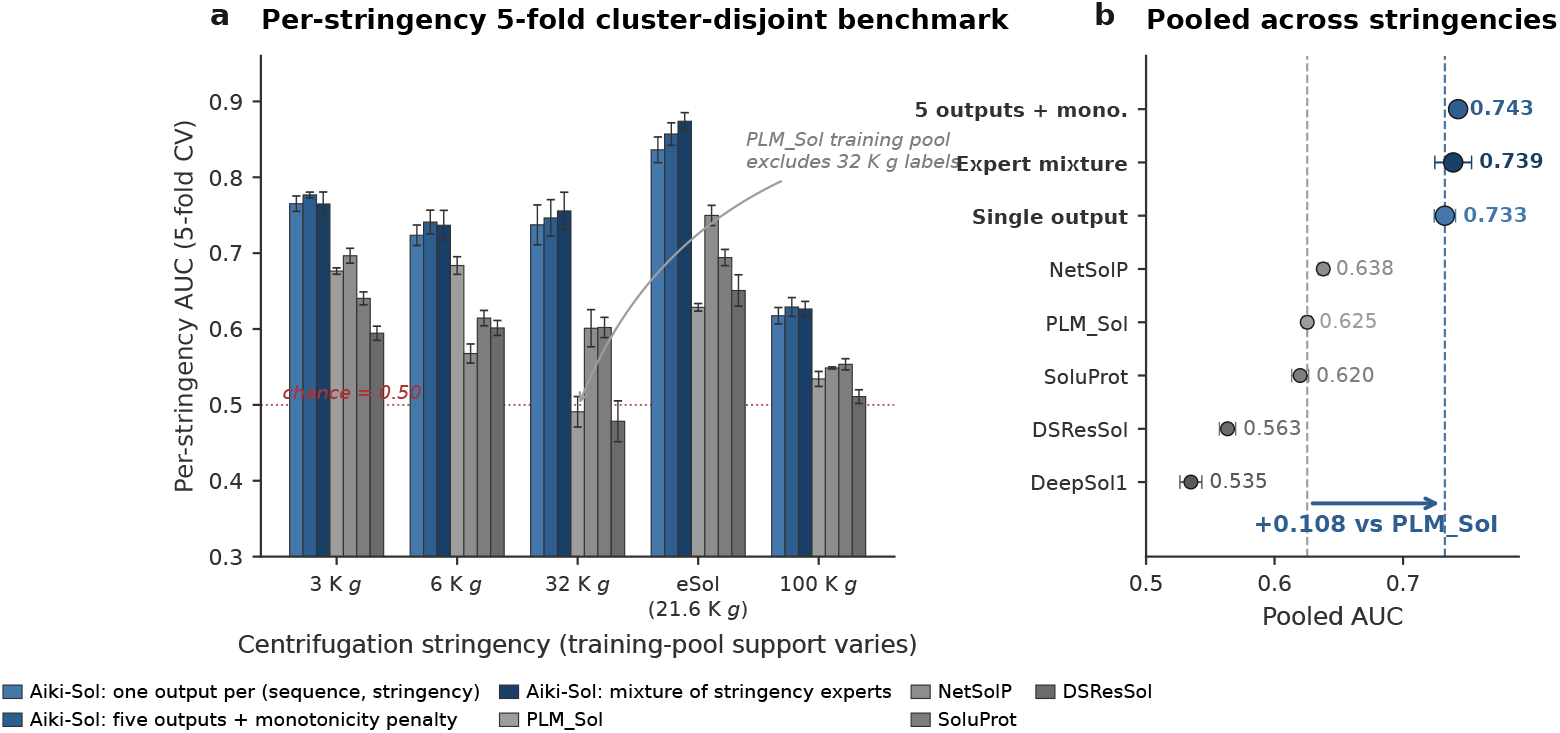
Cluster-disjoint per-stringency 5-fold benchmark under cluster-disjoint scoring. **(a)** Per-stringency ROC-AUC across all five centrifugation regimes for the three Aiki-Sol variants (5/5 folds each) and the two strongest published binary comparators (both at 5*/*5 folds). Error bars are sample standard deviation across folds. Dashed line at AUC = 0.5 marks chance performance. The 32,000 *g* stratum makes the per-protocol-vs-binary mechanism concrete: `PLM_Sol`’s training pool does not see this stringency and its 32,000 *g* AUC is 0.491 ± 0.020 (below chance, all five folds), while Aiki-Sol reaches 0.737–0.756 on the same proteins. **(b)** Pooled-across-stringencies AUC. Every Aiki-Sol variant lifts +0.108 to +0.118 over `PLM_Sol`’s 0.625 ± 0.003, an order of magnitude larger than the ±0.005–0.015 spread among Aiki-Sol’s own architectural variants.

#### The mechanism is exposed concretely on the 32,000 × *g* stratum

PLM_Sol’s training pool is the PSI:Biology subset and does not see 32,000 × *g* labels; its predicted probability anti-correlates with the binary outcome there (AUC 0.491 ± 0.020, below chance on all five folds), while Aiki-Sol’s 32,000 × *g* AUC across its variants is 0.737–0.756 on the same cluster-disjoint test proteins. `NetSolP` corroborates the pattern (32,000 × *g* AUC 0.601 ± 0.025). A binary predictor without per-protocol awareness can fail on a centrifugation stringency absent from its training pool without signalling to its user that the prediction is unreliable; a per-protocol-aware predictor delivers a calibrated prediction on all five stringencies simultaneously.

None of the three head-and-loss variants clears a pre-registered bar for declaring an improvement over the conditioned baseline (+0.020 pooled AUC with a bootstrap CI strictly positive on ≥4 of 5 folds; SI S4). How a per-regime threshold sets where the soluble/insoluble cut falls is the subject of §2.2.

### 2.2 Structure, capacity, and architecture exploration

The +0.108 lift of §2.1 dwarfs every head-and-loss elaboration on the same backbone. Three independent observations point to a simple mechanism: one shared protein ranking plus a per-regime threshold, rather than architecture, capacity, or structural information (Figure 3).

**Figure 3:**
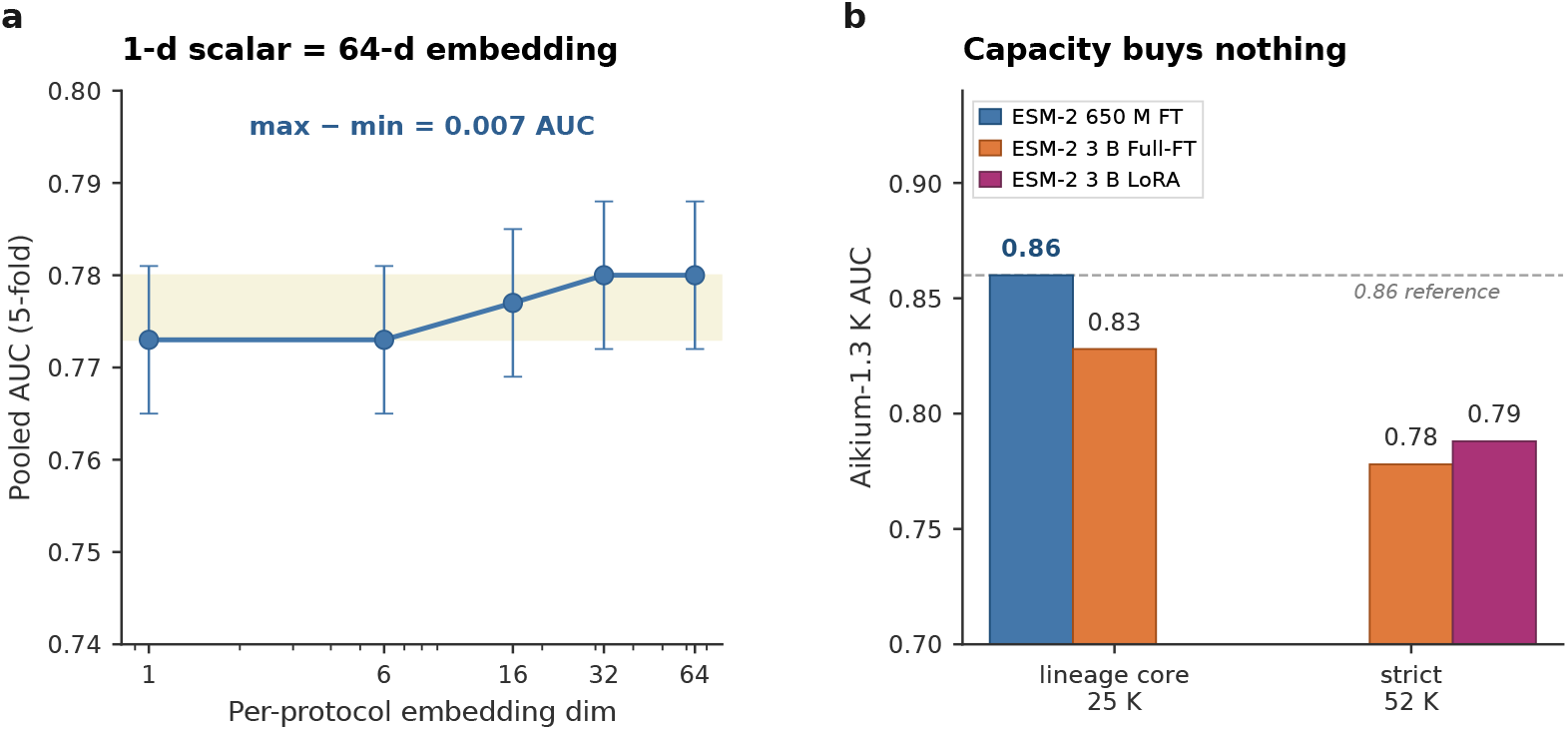
Architecture and capacity add no measurable signal beyond a single shared ranking. **(a)** A one-dimensional learned scalar per protocol (the conditioning channel reduced to a single bias vector indexed by stringency) matches a 64-dimensional learned embedding within ± 0.007 pooled AUC on the cluster-disjoint five-fold benchmark. The conditioning channel is consistent with a per-regime threshold on a shared ranking, not a protocol-specific protein representation. **(b)** Capacity-axis null on two license-clean training-pool scales: an ESM-2 3B Full-FT trails the conditioned 650M FT by ~ 0.04 AUC on the lineage-core 25K split, and 3B Full-FT and 3B LoRA on a license-clean 52K-row pool reach 0.778 and 0.788, with no lift over the smaller backbone. The structure-aware comparison (GATSol and ProtSolM scored on ESMFold-predicted structures for one benchmark fold) is in Table 1.

#### The mechanism reduces to one shared protein ranking plus a per-regime threshold

The five per-stringency predictions order proteins almost identically (mean pairwise Spearman *ρ* = 0.972 on the validation set, *N* = 3,102; *ρ* = 0.981 between the two most extreme regimes), so the model is essentially learning one shared score per protein and then asking, for each regime, where the soluble/insoluble cut should fall. **A one-dimensional learned offset per regime matches a** 64**-dimensional learned embedding within** ±0.007 **AUC** (Figure 3a); the conditioning channel encodes a per-regime threshold, not a learned protein representation. Removing it at training time drops scaffold AUC by ~ 0.24 at 650M; randomising the per-protein stringency identifier on a trained checkpoint drops stratified-mean AUC by 0.118. The five-element bias vector is the principal contributor.

#### Predicted structures do not lift performance on this benchmark

We predicted ESMFold v1 structures for fold 0 of the protocol-stratified pool (8,474 proteins) and scored the two published structure-aware tools, GATSol and ProtSolM, on them. Both land within the binary-comparator range (GATSol 0.613, ProtSolM 0.655 pooled; Table 1), below every language-model-only Aiki-Sol variant. On this cluster-disjoint per-stringency benchmark, supplying a predicted-structure channel does not beat the sequence-only frame. A complementary sequence-only check confirms the same conclusion from the other direction: per-stringency logistic regression on frozen `ESM-2` 650M masked-mean embeddings of the 84,809-protein pool reaches mean-of-per-stringency AUC 0.7534 ± 0.0049, within +0.003 of the strongest fine-tuned Aiki-Sol variant; the full fine-tune step contributes at most ~ 0.01 AUC beyond a frozen PLM with per-stringency awareness. A within-protein cross-stringency flip-prediction test (SI S4) locates the substantive learning in per-stringency calibration rather than the fine-tune step: per-stringency logistic regressions on frozen `ESM-2` embeddings clear a constant-bias floor by +13 pp, while fine-tuning the backbone adds only +2 pp on top (within bootstrap noise at *N* = 243 flipping pairs). What the fine-tune does sharpen is direction: it recovers 78% of the biophysically-expected flips while staying at chance on the suspicious ones, the likely source-label inversions a model should not reproduce.

#### Capacity adds no measurable signal at either license-clean pool scale tested (Figure 3b)

An `ESM-2` 3B Full-FT on the lineage-core 25K split trails the conditioned 650M FT by ~ 0.04 AUC on the prospective scaffolds; on a license-clean 52K-row pool, 3B Full-FT and 3B LoRA reach 0.778 and 0.788, with no lift over the smaller backbone. Capacity by itself does not unlock further lift across these scales; full training-pool details are in SI S4.

### 2.3 External cohorts and the cost of unannotated protocols

Five external cohorts (ProgSol-PsiBiology, ProgSol-NESG, ProtSolM-pdbsol, ProtSolM-DSResSol, ProtSolM-SoluProt) report a single binary “soluble” label; most lack the collection-protocol annotation needed to pick which of the five per-stringency Aiki-Sol probabilities applies. On these cohorts the useful comparison is no longer protocol-aware versus binary; it is how strongly each cohort’s training-pool overlap inflates apparent performance, and how each model scores on the cohort subset at measurably-zero overlap. Cluster-disjoint scoring at 25% identity shifts cohort-level AUCs by up to 16 pp on the highest-overlap cohort (PSI:Biology, 79% training-pool overlap) but by less than 0.06 AUC on the remaining four (SI S5).

#### The canonical-147K-trained deployment checkpoint lifts cohort-mean AUC to 0.825 across five external cohorts

Under matched protocol-aware aggregation (Table S5), Aiki-Sol reaches 0.803 ± 0.005 across the cluster-disjoint 5-fold partition of canonical-147K and 0.825 for the full-pool deployment checkpoint, lifting both `NetSolP` (0.701 cohort-mean) and `PLM_Sol` (0.637) by +0.10–0.19 AUC. The cohort-by-cohort training-pool overlap of canonical-147K is asymmetric and explicitly measured: 0% exact-sequence overlap with all three ProtSolM-derived cohorts (pdbsol, DSResSol, SoluProt; the canonical pool excludes the ProtSolM-license family by construction) and 46% / 15% exact-sequence overlap with ProgSol-PsiBiology / ProgSol-NESG (source corpora that contribute to canonical-147K). The clean three-cohort subset is the defensible zero-leakage comparison: on it Aiki-Sol reaches 0.754 (5-fold mean) and 0.772 (full-pool deployment), versus `NetSolP` 0.613 and `PLM_Sol` 0.676. The gap is a +0.10 to +0.16 AUC lift under measurably-zero training-pool overlap (NetSolP and PLM_Sol training corpora do not contain ProtSolM-family proteins, so the overlap measurement is symmetric on this comparison). The two leaky cohorts (PSI:Biology, NESG) reflect partial memorisation on Aiki-Sol’s side and would be expected to drop under cluster-disjoint re-scoring against canonical-147K; the protocol-stratified 5-fold benchmark of §2.1 and the canonical-147K parallel in Table S6 remain the leakage-free headlines. Per-protein predictions and per-cohort overlap counts are released alongside the paper.

#### Cohort-mean lift decomposes by overlap

The full-pool deployment gains +0.018 on the zero-leakage subset (0.754 → 0.772), the genuine benefit of the roughly 20% larger training pool, plus an overlap-correlated +0.039/+0.022 on PSI:Biology/NESG (46% and 15% overlap respectively), the memorisation effect the cluster-disjoint 5-fold checkpoints are designed to expose.

#### Below-chance binary failure recurs on canonical-147K

`PLM_Sol` falls below chance on 32,000*g* (0.491 ± 0.020) on the 85K benchmark; `NetSolP` falls below chance on 6,000*g* (0.450 ± 0.029) on canonical-147K (Table S6). Which stringency falls below chance shifts with dataset composition, confirming that a single binary predictor cannot maintain calibration across stringencies.

A small Aikium-internal cohort of 1,261 designed recombinant scaffolds (binary inclusion-body vs. cytosolic SDS-PAGE classification, with ID-thresholded labels and high tagged-input construct heterogeneity) is held out at 0% exact-sequence overlap with the released training pools and used only as a deployment-target description (SI S6, label-construction caveats). The released checkpoint reaches AUC in the high 0.8s on this cohort.

## 3 Discussion

### The centrifugation regime is a latent confound in *E. coli* solubility datasets that the field has been collapsing into a single label

The protein’s underlying state does not change between regimes; what changes is which lysate subpopulation is counted as “soluble” and which is pelleted. For nearly a decade, sequence-based predictors of recombinant solubility have collapsed this confound into a single label and been tested on partitions that share substantial sequence identity with their training pools. The overlap inflates the reported numbers directly; collapsing the confound is quieter but lets a protocol-specific result read as a general claim about the protein. The two compound: a model that has seen a near-duplicate can reproduce its protocol-specific label without having learned anything transferable. The strongest published binary comparator on our protocol-stratified cluster-disjoint partitions falls *below chance* on the 32,000 × *g* stratum (AUC = 0.491 ± 0.020, all five folds; `PLM_Sol`’s training pool does not see this stringency); a second binary comparator (DSResSol; AUC 0.478 at the same stratum) reproduces the pattern on a different training set. A binary predictor that has no per-protocol awareness can fail on a stringency absent from its training pool *without signalling that the prediction is unreliable*. Under cluster-disjoint scoring on a single benchmark, no published binary comparator from 2018–2025 pools above 0.66; reported gains over four years [19, 20] (~ 0.04 pooled AUC) are of the same magnitude as the cluster-disjoint inflation correction (~ 0.06 on three of five public cohorts; SI Table S4), making real algorithmic improvements difficult to disentangle from evaluation-protocol effects without cluster-disjoint comparison.

### Architecture, capacity, and predicted structure add no measurable signal

Three independent observations (§2.2) point in the same direction: structure-aware predictors, scored on ESMFold structures for the benchmark, do not beat the language-model-only frame; head-and-loss variants on top of the conditioned backbone clear neither a +0.020 AUC threshold nor a paired-bootstrap CI test; capacity above 650M parameters does not clear the smaller backbone at either pool scale tested. The data are consistent with one shared protein ranking (per-regime predictions agree at Spearman *ρ* = 0.972) plus a per-regime threshold; a one-dimensional learned offset per regime matches a 64-dimensional learned embedding within ± 0.007 AUC on this benchmark. A fine-tuned ESM-2 650M backbone with a five-element bias vector therefore reaches the headline performance. For the protein scientist at deployment, the practical consequence is that the relevant solubility prediction is no longer a single number but the one matched to the centrifugation regime their downstream workflow uses (or the marginal head when the workflow is unspecified). The centrifugation regime, implicit in existing solubility datasets, is now an explicit parameter the user selects.

### Limitations and open directions

“Solubility” here is the fraction of expressed protein recovered in the supernatant after centrifugation, or the equivalent SDS-PAGE band. It is an *E. coli* recombinant-expression readout, distinct from the colloidal or formulation stability optimised in downstream development. Aiki-Sol is sequence-only, trained on *E. coli* BL21(DE3), and scores the whole protein; it does not resolve individual point mutations (mean Spearman *ρ* = 0.07 on the SoluProtMutDB deep-mutational panel; SI S6). The binary frame is also near its ceiling as currently collected: contributing sources disagree on the label for 53% of multiply-measured proteins, and once protein length, hydrophobicity, and net charge are accounted for, the laboratory a measurement came from carries essentially no further information about the label (SI S2). Two routes could move past this. On the structure side, ESMFold-predicted coordinates add little that the sequence representation does not already carry [19], and several structure-aware predictors [15, 16, 21, 22] could not be run on the cluster-disjoint partitions at all (closed-source or web-only dependencies); a real test of the structural axis would need experimentally determined coordinates. On the data side, recording the centrifugation protocol at the point of measurement, rather than reconstructing it post-hoc from heterogeneous sources [20], would let the field measure directly the variable this paper has had to infer.

## 4 Methods

### 4.1 Dataset and benchmark

Four nested data pools support the analyses in this paper (Table 2). Two are the training pools of the released models; the other two are benchmark and scale-probe pools.

The protocol-stratified 84,809-protein pool annotates each protein with its centrifugation stringency and is 100% known-stringency by construction; it is the benchmark pool of §2.1 and §2.2. The canonical-147K pool extends it with binary-unknown-stringency proteins and continuous mg/mL eSol measurements (license-restricted sources excluded) and is the training pool of the released Aiki-Sol deployment checkpoint (§4.6; Apache 2.0). The research-tier 229K pool additionally includes CC-BY-NC-ND sources and supports a research-tier Aiki-Sol-research checkpoint; its in-distribution evaluation on ProtSolM-derived external cohorts is reported separately as memorisation rather than generalisation.

All pools are partitioned into cluster-disjoint 5-folds at 25% sequence identity and 80% bidirectional coverage using MMseqs2 [10]. Binary labels across pools collapse three operationally distinct phenotypes [24, 25] that we treat as a unified label for tractability, with the resulting label noise characterised in SI S2.

#### Per-stringency positive rates are non-monotonic with *g*-force

(full breakdown SI S1). This is a fingerprint of the upstream-source mixture (different consortia operationalise different *g*-force cuts with different positive-rate biases) rather than a biophysical property of *g*-force itself. We do not balance positives against negatives within each centrifugation regime during training; this preserves deployment-time interpretability of *P* (soluble | *g*) as inheriting the upstream-source prior. The loss instead carries two correction terms against per-stringency skew (Table S1, §4.6): a cross-supervision branch anchoring the mean of the five sigmoids to each protein’s binary label, and a squared-ReLU penalty on non-monotonic adjacent-stringency pairs.

### 4.2 Cluster-disjoint cross-predictor benchmarking

For cross-predictor comparison on a public cohort, we cluster the union of (i) the cohort’s protein set and (ii) every training pool known to have been used by the predictors under comparison (PSI:Biology, eSol/PURE, ProtSolM training, the `NetSolP` training set, the `PLM_Sol` training set, and the Aiki-Sol Dataset splits) at 25% identity with ≥ 80% coverage, and retain only test proteins whose cluster does not intersect any training pool. The cluster assignments are released as machine-readable artefacts so any future predictor can be benchmarked against the same partitions.

### 4.3 Protocol-stratified five-fold cross-validation

The protocol-stratified 84,809-protein pool is clustered at 25% identity with ≥ 80% bidirectional coverage and partitioned into five folds with every cluster entirely in one fold. A stratification rule requires every (stringency, primary-center) stratum with ≥ 30 proteins pool-wide to be represented by ≥ 3 proteins in each fold’s test partition, preventing single-stringency inflation. The headline metric is per-stringency AUC mean ± standard deviation across folds on each fold’s cluster-disjoint test partition.

### 4.4 Architecture

All Aiki-Sol variants share a fine-tuned `ESM-2` 650M backbone [2] (end-to-end, layer-wise learning-rate decay), masked-mean pooling over residues, and a soft ordinal-monotonicity penalty (Σ ReLU(Δ*p*)^2^ over adjacent stringency pairs, weight 0.2; Table S1 branch 4) that enforces tier monotonicity; they differ only in the head and in how the centrifugation regime conditions the prediction. The released deployment checkpoint (§4.6) emits five per-regime probabilities plus a marginal output directly from a linear head and takes no regime input. The conditioned-baseline variant of §2.1 instead takes the regime as a categorical input: the pooled representation is concatenated with a 32-dimensional learned regime embedding and passed through a three-layer MLP with ReLU and dropout 0.1, and one checkpoint is then scored at any regime by indexing the embedding table. An embedding-dimension sweep across *{*1, 6, 16, 32, 64*}* moves pooled AUC by only 0.007 (Figure 3a).

### 4.5 Tag-aware sequence normalisation

Recombinant proteins in solubility datasets carry hexahistidine (H_6_), HiBit (VSGWRLFKKIS, often flanked by (GGGGS)_n_ linkers), and an N-terminal methionine that may or may not have been cleaved. Public predictors disagree on whether to strip these and published evaluations rarely report the choice; the omission moves headline AUC by 0.01–0.05 (SI S3). We canonicalise every sequence the model sees, at both training and inference, through one entry point `normalize_sequence`: H_6_ runs are detected at either terminus and stripped within a windowed flanking region; HiBit and its (GGGGS)_n_ linkers are removed; an optional N-terminal methionine is handled configurably (canonical: do not force Met-cleavage at scoring time). A post-normalisation invariant is asserted (retained-H_6_ and retained-HiBit rates both *<* 5%) and the score is rejected otherwise. All numbers in this paper, for our model and every published comparator, are produced under this canonical normalisation.

### 4.6 Training

#### Optimiser and schedule

All Aiki-Sol variants share AdamW (weight decay 0.01, backbone lr 5 × 10^−5^, head lr 5 × 10^−4^) with layer-wise learning-rate decay 0.85 [9], gradient checkpointing under `bfloat16` autocast, effective batch 32, sequence length ≤ 1,022. Validation is monitor-only and never used for checkpoint selection; epoch count is pre-registered per training pool. The four Aiki-Sol variants in Table 1 all train for two epochs and differ only in the head and loss specification; the per-variant loss forms are in SI S4.

#### Released checkpoint (Aiki-Sol)

The deployment artefact is one fine-tuned ESM-2 650M backbone, masked-mean pooled, projected through Linear(1280, 256) → ReLU → Linear(256, 6) to six outputs: *p*_0_–*p*_4_ are per-stringency soluble probabilities at 3,000*g*/10 min, 6,000*g*, 32,000*g*, eSol/21,600*g*, 100,000*g*; *p*_5_ is a dedicated marginal head trained to predict *P* (soluble | sequence) without stringency conditioning, supervising the canonical-147K proteins whose source record does not recover the centrifugation stringency. Three annotation classes route each training sample to a subset of seven loss branches (Table S1): known-stringency binary samples supervise the matching *p*_*g*_ and *p*_5_; unknown-stringency binary samples supervise *p*_5_ directly and cross-supervise 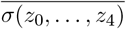 at weight 0.5; continuous eSol/PURE measurements supervise *σ*(*z*_3_) and *σ*(*z*_5_) under MSE. A soft ordinal-monotonicity penalty (weight 0.2, Σ ReLU(Δ*p*)^2^ over the (*p*_0_, *p*_1_), (*p*_1_, *p*_2_), (*p*_2_, *p*_4_) pairs) applies to every training sample.

#### Schedule

Aiki-Sol (canonical-147K, Apache-tier) trains for two epochs; Aiki-Sol-research (research-tier 229K, CC-BY-NC-ND) for three epochs under an otherwise identical recipe. The deployment full-pool checkpoint uses a 95%*/*5% train/val split; per-fold checkpoints trained on the cluster-disjoint 5-fold partition produce Table S6. Both checkpoint families are released. For frozen-PLM baselines (`ESM-C` 300M, ESM-C 600M [26], `ProtT5-XL`-XL) we train a conditioned linear head over masked-mean-pooled representations [18].

#### Holdout-pool overlap reporting

Every holdout AUC reported carries its measured exact-sequence overlap with the run’s training pool, distinguishing *in-distribution* (≥ 1%) from *held-out* (*<* 1%); per-cohort overlap figures are reported in SI S3.

## Data availability

The canonical 147K-row Apache-tier training pool of Aiki-Sol, its 5-fold cluster-disjoint partition assignments at 25% identity, the Apache-tier deployment checkpoint, the five per-fold checkpoints, the per-cohort prediction CSVs, and the aggregated result JSONs of the released checkpoints are deposited at Zenodo: 10.5281/zenodo.20151817 (concept DOI; resolves to the latest published version). The training pool, splits, and predictions are released under CC BY 4.0; the Apache-tier deployment checkpoint and per-fold checkpoints under Apache 2.0.

The research-tier Aiki-Sol-research checkpoint and its per-cohort predictions are released at the same Zenodo deposit under CC-BY-NC-ND 4.0, inheriting the most-restrictive upstream-source licence tier. The underlying 229,349-row training CSV extends the canonical pool with rows under CC-BY-NC-ND 4.0 (SoluProtMutDB and the ProtSolM family) and Springer Nature defaults for additional sources; it is NOT redistributed verbatim. The deposit’s per-source manifest (`research_tier/n3v2_source_manifest.csv`) names each upstream source so researchers can re-fetch from the original distributors and rebuild the training pool under their own institutional licensing.

The 84,809-protein protocol-stratified benchmark pool of §2.1 mixes Apache- and CC-BY-NC-tier proteins from the upstream sources documented in SI §S1 (TargetTrack, eSol/PURE, seven structural-genomics consortia); the pool itself is not redistributed verbatim, and the canonical 147K-protein pool deposited above is its largest fully-redistributable subset. The in-house held-out evaluation cohorts (two engineered-scaffold datasets and three nanobody datasets) used in this work for evaluation only are not in any released training pool and are not redistributed.

## Code availability

Source code for data assembly, split construction, model training, inference, and figure reproduction is publicly available under an Apache 2.0 license at https://github.com/aikium-public/aiki-sol and archived at the same Zenodo DOI as the dataset deposit above. The deployment artefact (Aiki-Sol, §4.6) is distributed as a Python package with the same license and a `predict(seq)` entry point (`pip install aikisol`); Apache 2.0-licensed model weights are fetched on first call from the Zenodo deposit. Source code and pretrained weights for the Aiki-Sol-research checkpoint (trained on the research-tier 229K pool, which includes upstream sources under CC-BY-NC-ND 4.0 that prohibit redistribution-as-weights of the trained model) are released at the same repository under a separate research-use notice that flags those upstream license restrictions.

## Acknowledgements

We thank the NESG, PSI:Biology, MCSG, CSGID, and NYSGRC consortia for making their structural-genomics data publicly available through TargetTrack, and the curators of Target-Track itself for maintaining the open access to consortium-level expression and solubility records that this work builds on. We thank the eSol/PURE team [4] for the cell-free aggregation propensity dataset that anchors the eSol stratum of our protocol-aware benchmark. We thank the developers of the protein language models evaluated in this work (ESM-2 and ESM Cambrian from Meta and EvolutionaryScale, and ProtT5-XL from TUM/Rostlab) for open model weights, and the MMseqs2 [10] team for the sequence-clustering tool on which our cluster-disjoint partitions depend. This work was supported by Aikium Inc.

## Author contributions

R.R. curated the initial version of the solubility dataset, trained models that recapitulated the leading non-stringency based models, and helped write the final manuscript. R.S.M. contributed with technical feedback and manuscript editing. S.S. provided the protein sciences domain knowledge for the centrifugation-protocol interpretation and contributed to manuscript revision. V.M. conceived the project, curated the final dataset, implemented the stringency-aware architectural variants and benchmarks, performed the experiments, compiled the results, and wrote the manuscript.

## Supplementary Information

### S1 Dataset construction

The strict-curation lineage core of Aiki-Sol Dataset contains 35,808 *E. coli* proteins drawn from TargetTrack records [6] cross-matched against the eSol/PURE collection [4] and seven structural-genomics consortia. Each protein retains its centrifugation speed. Per-center centrifugation parameters are assigned from primary TargetTrack `protocolText` extractions: a manual review of the documented protocols places NESG, NYSGRC, and CESG at HIGH confidence (rotor or g-force explicit in the source XML) and MCSG, CSGID, and EFI at MEDIUM confidence (rotor inferred from rpm and protocol context). The corrected mapping is released as `h6_v12_center_lookup.csv` at the same Zenodo DOI as the dataset (§4.6); seven additional PSI:Biology centers (NYSGXRC, NYCOMPS, SECSG, BSGC, TB, BSGI, OCSP) and four further centers (JCSG, RSGI, SGX, SSGCID) lack a documented soluble-call step and are released as “unknown_g” for transparency.

#### Per-stringency breakdown of the lineage core

(*N* = 35,808): 3,000*g* NESG/PSI 12,617 (65% positive), 6,000*g* NYSGRC 740 (99%), 32,000*g* MCSG/others 8,009 (54%), 100,000*g* CSGID 1,314 (99.7%), 21,600*g* eSol/PURE 5,708 (45%), and Unknown *g* 7,420 (65%). The documented-protocol review (Methods §4.1; released artifact `h6_v12_center_lookup.csv`) corrects MCSG to the 100,000*g* class (32,000 rpm × 40 min ultracentrifugation; rotor inferred) and CSGID to the 32,000*g* class (SS-34 @ 17,355 rpm × 1 hr *≈* 37,000 × *g* per protocolText), shifting ~ 5,500 proteins between strata; headline §2.1 numbers are independent of which centre supplies which stratum.

The corpus is partitioned into cluster-disjoint train (25,026), validation (3,102), test (3,105), and external (4,486) splits at 25% identity using MMseqs2 [10]. The lineage core is a strict-curation subset of the broader curated literature collection.

### S2 Label-noise floor characterisation

Four converging diagnostics bound what binary classification can deliver in this domain as currently posed. (i) Inter-source label disagreement: on a 135,268-protein union with 7 sources and 18,199 multi-source clusters, 53% of multi-source clusters have at least one source disagreeing on the binary label; concordant-cluster AUC reaches 0.904, discordant-cluster AUC 0.697. A finer-grained re-aggregation of the same disagreements across canonical centrifugation stringencies finds that of the 13,224 proteins flagged as cross-source-disagreeing in the protocol-stratified pool, only 209 (1.6%) are true *same-stringency* cross-source binary disagreements at known stringencies; the remainder are cross-stringency contrasts (which biophysics expects to flip across protocols) or disagreements involving unknown-stringency observations. The per-stringency pool is therefore much cleaner than the raw union-level disagreement count suggests once the centrifugation regime is controlled. (ii) Source-as-random-effect intraclass correlation: logistic mixed-effects model with source as random intercept and sequence-derived covariates (length, GRAVY, charge per residue) as fixed effects gives ICC_source_ = 0.000 versus 0.033 uncontrolled one-way ANOVA: source identity is conditionally independent of label given sequence. (iii) Architectural saturation: seven 650M-parameter variants cluster within a 0.82–0.86 ROC-AUC band on the prospective scaffold set under unified scoring. (iv) Field-wide NESG band: across eight PLM-based predictors over four years, NESG independent-test AUC has moved by approximately 0.04 [19, 20]; the same magnitude as the cluster-disjoint inflation correction in Table S4.

### S3 Per-cohort training-pool overlap

#### In-distribution versus held-out

In any real-world deployment both familiar (in-distribution) and novel (out-of-distribution) protein sequences appear; both matter, and conflating them inside one AUC obscures which deployment scenario each number addresses. Each holdout AUC reported in this paper carries its measured normalised sequence-identity overlap with the specific training pool used for the run that produced it. We distinguish two zones: *in-distribution* (≥ 1% exact-match overlap, capturing deployment behaviour on proteins similar to the training corpus) and *held-out* (*<* 1% exact-match overlap, capturing generalisation to novel sequences). Both zones are reported and never collapsed.

The protocol-stratified 84.8K pool is the training pool against which the architectural verdicts of §2.1 are computed; both the prospective scaffold cohort and the cluster-disjoint per-fold test partition are at 0% exact-sequence overlap with it. Every released checkpoint’s training pool is likewise at 0% exact-sequence overlap with the prospective scaffold cohort (§S6).

#### Per-cohort training-pool overlap with Aiki-Sol

Cohort-by-cohort training-pool overlap is also the dominant driver of the cross-predictor ranking in §S5. The five external cohorts and their measured exact-sequence overlap with Aiki-Sol’s training set are: ProgSol-PsiBiology 79%; ProgSol-NESG 32%; ProtSolM-pdbsol 24%; ProtSolM-DSResSol 23%; ProtSolM-SoluProt 18%. Predictors trained on overlapping pools are subject to the same magnitude of inflation when scored on the same union splits. `NetSolP` (trained on PSI:Biology) is in-distribution to PsiBiology and NESG cohorts; `PLM_Sol` (trained on a UniRef-derived pool covering PDB-aligned sequences) is in-distribution to pdbsol; the corrected leaderboard reflects each predictor’s training-distribution match more closely than any absolute architectural advantage.

### S4 Supplementary architectural diagnostics

Three further architectural diagnostics support the noise-floor verdict of §S2 and the saturation framing of §S5.

**Figure S1:**
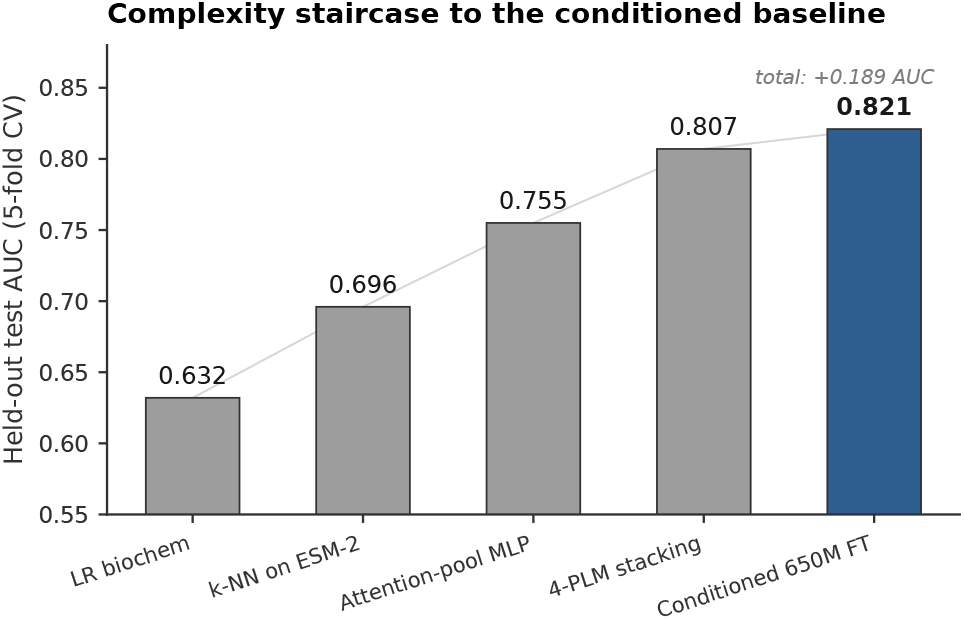
Complexity staircase. Held-out test AUC (5-fold CV) as model complexity grows from a length-and-composition logistic regression (0.632) through k-NN on ESM-2 650M embeddings (0.696), an attention-pooled MLP (0.755), and four-PLM stacking (0.807), to the conditioned ESM-2 650M fine-tune (0.821). Within the fine-tuned-PLM frame the marginal gain from further architectural elaboration is small; the full architecture and training recipe are in Methods §4.6.

#### Aiki-Sol loss branches

Table S1 expands the deployment-checkpoint loss specification of §4.6. Each training sample activates a subset of seven branches based on its annotation class (`binary_at_known_strg`, `binary_unknown_strg`, `continuous_eSol`); the ordinalmonotonicity penalty applies to every sample. On a typical mini-batch every sample contributes to the monotonicity penalty and to one or more BCE/MSE terms; no sample contributes to a loss branch that conflicts with its label. Proteins whose source centrifugation protocol is systematically biased against the corpus-level convention are dropped at preprocessing rather than fed at a conflicting label value.

**Table S1:** Aiki-Sol loss branches.

| Branch | Active on samples where annotation = | Weight | Loss form | Target |
| --- | --- | --- | --- | --- |
| 0a | `binary_at_known_strg` | 1.0 | BCE | $\sigma(z_g)$ vs binary label |
| 0b | `binary_at_known_strg` | 1.0 | BCE | $\sigma(z_5)$ vs same binary label |
| 1a | `binary_unknown_strg` | 1.0 | BCE | $\sigma(z_5)$ vs binary label |
| 1b | `binary_unknown_strg` | 0.5 | BCE | $\sigma(z_0, \dots, z_4)$ vs same label |
| 3a | `continuous_eSol` | 1.0 | MSE | $\sigma(z_3)$ vs normalised mg/mL |
| 3b | `continuous_eSol` | 1.0 | MSE | $\sigma(z_5)$ vs same target |
| 4 | all samples | 0.2 | $\sum \text{ReLU}(\Delta p)^2$ | monotonicity over $(p_0, p_1)$, $(p_1, p_2)$, $(p_2, p_4)$ |

#### Within-protein cross-stringency flip test

A stricter question than per-stringency AUC is whether a model recovers *within-protein* solubility changes across centrifugation regimes. We assembled every protein in the curated corpus carrying *≥* 2 known-stringency binary observations whose label *disagrees* across stringencies (the same protein scored soluble at one regime and insoluble at another), giving 240 proteins and 243 flipping pairs. A pair is *expected* when the gentler stringency is the soluble one (152 pairs; biophysically consistent, since more centrifugal force pellets more material) and *suspicious* when the harsher stringency is (91 pairs; not biophysically expected, and most likely a source-label inversion). The test asks whether a model predicts the direction of the flip. We compare three models (Table S2): a constant-bias floor (one unified logistic regression plus a per-stringency log-odds offset, which carries zero within-protein cross-stringency information by construction); five independent per-stringency logistic regressions on frozen ESM-2 650M embeddings; and the fine-tuned Aiki-Sol architecture, scored cluster-disjoint with each protein evaluated by the fold checkpoint that did not train on it.

The frozen-embedding regressions clear the constant-bias floor by +13 pp, and fine-tuning the backbone adds only a further +2 pp, within bootstrap noise at *N* = 243: the within-protein cross-stringency signal is already present in the frozen representation, and the per-stringency calibration channel, not the fine-tune, is the load-bearing operation. The fine-tune does sharpen one thing, direction: Aiki-Sol recovers 78% of the biophysically-expected flips while sitting at chance on the suspicious flips, which is where a model that has learned solubility rather than memorised source labels should sit. At this substrate size (243 pairs) the test is a mechanistic diagnostic, not a high-powered statistical contrast.

**Table S2:** Within-protein cross-stringency flip-direction recovery: the fraction of flipping pairs whose predicted direction matches the label direction. Three models on the 243-pair flip substrate; Aiki-Sol is scored cluster-disjoint via held-out fold checkpoints.

| Model | Overall<br>($n = 243$) | Expected<br>($n = 152$) | Suspicious<br>($n = 91$) |
| --- | --- | --- | --- |
| Constant-bias floor | 53.9% | 56.6% | 49.5% |
| Frozen ESM-2 650M + per-stringency LR | 67.1% | 69.7% | 62.6% |
| Aiki-Sol (fine-tuned, cluster-disjoint) | **69.1%** | **78.3%** | 53.8% |

#### Per-stringency training-pool prior vs. Aiki-Sol learned offsets

Table S3 lays the training-pool positive rate, the Aiki-Sol 5-fold per-stringency test AUC (Methods §4.3), and the population mean prediction on the cross-stringency flip-protein subset defined above side-by-side. The model’s learned per-stringency offset tracks the training prior, not a g-force monotone.

**Table S3:**
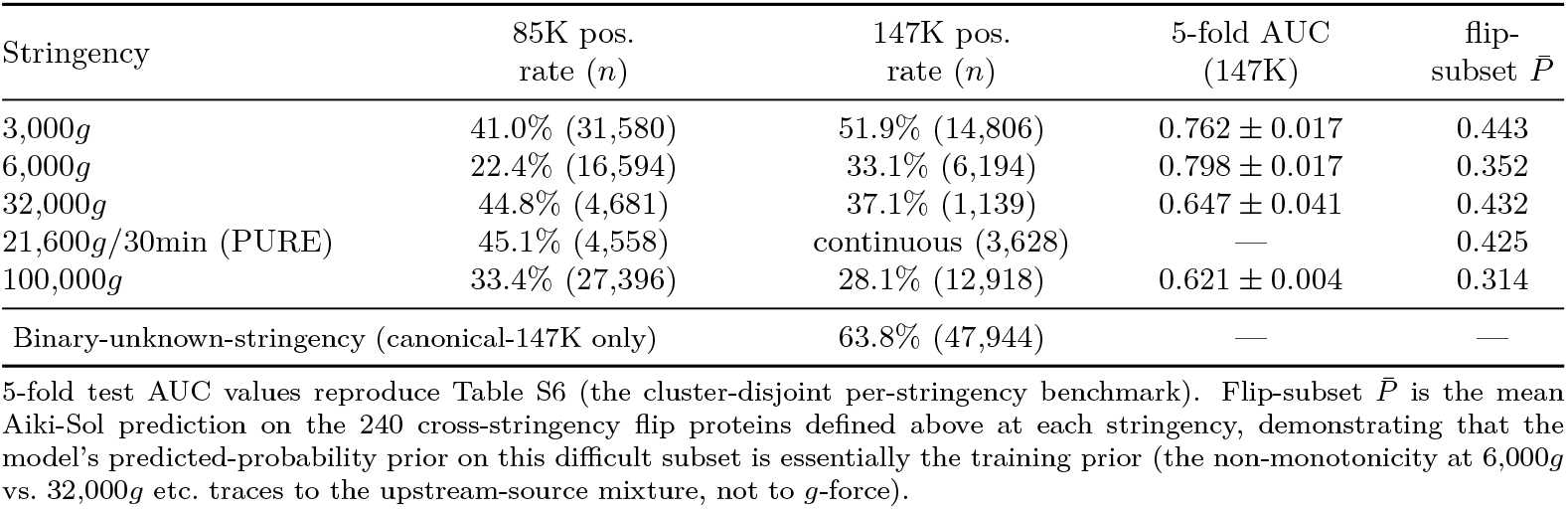
Per-stringency training-pool positive rate (the model’s calibration prior), Aiki-Sol 5-fold test-partition AUC on each stringency, and the population-mean Aiki-Sol prediction on the within-protein cross-stringency flip-protein subset defined above (*N* = 240 proteins). Both training pools are non-monotonic with *g*-force, and the model’s learned offsets inherit this non-monotonicity. The eSol stratum on canonical-147K is routed to a continuous regression target, so a per-stringency binary AUC is not reported on canonical-147K for that cell.

| Stringency | 85K pos.<br>rate ($n$) | 147K pos.<br>rate ($n$) | 5-fold AUC<br>(147K) | flip-<br>subset $\bar{P}$ |
| --- | --- | --- | --- | --- |
| 3,000*g* | 41.0% (31,580) | 51.9% (14,806) | $0.762 \pm 0.017$ | 0.443 |
| 6,000*g* | 22.4% (16,594) | 33.1% (6,194) | $0.798 \pm 0.017$ | 0.352 |
| 32,000*g* | 44.8% (4,681) | 37.1% (1,139) | $0.647 \pm 0.041$ | 0.432 |
| 21,600*g*/30min (PURE) | 45.1% (4,558) | continuous (3,628) | — | 0.425 |
| 100,000*g* | 33.4% (27,396) | 28.1% (12,918) | $0.621 \pm 0.004$ | 0.314 |
| Binary-unknown-stringency (canonical-147K only) |  | 63.8% (47,944) | — | — |

### S5 External cohorts: cluster-disjoint benchmark

The five external cohorts (ProgSol-PsiBiology, ProgSol-NESG, ProtSolM-pdbsol, ProtSolM-DSResSol, ProtSolM-SoluProt) report a single binary “soluble” label without the protocol-of-collection annotation. We benchmark predictors on these cohorts at cluster-disjoint partitions (25% identity); for Aiki-Sol we report the released deployment checkpoint (§4.6) at protocol-aware aggregation (cohort-specific stringency-output averaging per documented collection protocol; details and per-cohort assignments in the footnote to Table S5). Cluster-disjoint scoring shifts cohort-level AUCs by up to 16 pp on the highest-overlap cohort (PSI:Biology, 79% overlap) and by |Δ| ≤ 0.06 on the other four (Table S4, two of which are within ±0.01 of union-split because the cluster-safe filter happens to retain a comparably-difficult subset); under the corrected partitions the cross-predictor ranking (Table S5) differs sharply from published reports.

#### Canonical-147K parallel benchmark

Table S6 replicates the per-stringency comparison of §2.1 on the larger curated canonical-147K pool (the training pool of the Aiki-Sol deployment checkpoint; architecture and training recipe in §4.6). The five Aiki-Sol fold checkpoints are scored on the canonical-147K cluster-disjoint test partition (34,990 known-stringency proteins across 5 folds; partition built by sequence clustering at 25% identity, 80% coverage on the 174K-protein canonical pool). The comparator rows of this table are not new scoring runs: they re-bin the existing per-protein predictions from §2.1’s protocol-stratified-85K benchmark under the canonical-147K 5-fold partition assignment (the 34,990 known-stringency test proteins of canonical-147K are a strict subset of protocol-stratified-85K by construction; coverage is 100% for 4 of 5 tools and 99.99% for SoluProt due to the same 3 mmseqs2 alignment failures documented in footnote a of §2.1). The protocol-aware lift survives the benchmark change: Aiki-Sol cohort-mean 0.7069 ± 0.0149 vs. best comparator (`PLM_Sol` 0.6400 ± 0.0959) gives +0.067 AUC, of the same order of magnitude as the +0.108 on protocol-stratified-85K from §2.1.

**Table S4:** Aiki-Sol AUC under union-split versus cluster-disjoint scoring across five external cohorts, both at the protocol-aware aggregation of Table S5. The inflation effect is concentrated on the cohort with the largest training-pool overlap (PSI:Biology); on cohorts with *<* 35% overlap the union-split and cluster-disjoint AUCs differ by |Δ| ≤ 0.06, and for two cohorts (NESG, pdbsol) cluster-disjoint scoring is in fact within ±0.01 of union-split (the cluster-safe filter happens to retain the harder-to-distinguish subset). The honest generalisation: training-pool overlap inflation is real but heavily cohort-specific, not a uniform 5–14 pp correction.

| Cohort | Union-split | Cluster-disjoint | $\Delta$ | Train overlap<sup>a</sup> |
| --- | --- | --- | --- | --- |
| ProgSol PsiBiology | 0.925 | 0.764 | −0.161 | 79% |
| ProgSol NESG | 0.776 | 0.786 | +0.010 | 32% |
| ProtSolM pdbsol | 0.664 | 0.667 | +0.004 | 24% |
| ProtSolM dsressol | 0.715 | 0.658 | −0.057 | 23% |
| ProtSolM soluprot | 0.691 | 0.631 | −0.060 | 18% |

<sup>a</sup> “Train overlap” is the fraction of cohort proteins sharing a sequence cluster (25% identity, 80% bidirectional coverage) with any lineage-core training-pool protein. Exact-sequence overlap is substantially higher (PSI:Biology 99.5%, NESG 97.0%; §S3); the cluster-disjoint 25%-identity filter retains only proteins with no near-homolog in the training pool.

The four-year, eight-predictor NESG drift band of ~ 0.04 AUC [19, 20] sits within the same magnitude as the cluster-disjoint inflation correction; the reported gains likely reflect both real algorithmic improvements and evaluation-protocol effects whose relative magnitudes are difficult to disentangle without cluster-disjoint comparison.

### S6 Prospective recombinant scaffold deployment cohort

#### What this cohort is and is not

The prospective evaluation uses 1,261 designed recombinant scaffolds (Aikium-internal constructs; ~ 1,000 from the inclusion-body-vs-cytosolic SDS-PAGE classification effort plus a small panel of additional scaffolds and positive controls). The cohort is reported here as a *deployment-target description*, not a comparator-axis benchmark. Two label-construction caveats apply, both inherited from the way the underlying assay was run rather than from the analysis pipeline, and they preclude any rank-order interpretation across architectures on this cohort. *(i) ID-thresholded labelling*. The binary labels follow a stair-step pattern keyed to construct ID rather than to a sequence-level assignment: IDs 1–524 are all classified 0, IDs 625+ are all classified 1, with a ~ 100-row mixed transition zone in between. This is a consequence of campaign-level batch labelling during construct prioritisation, not a per-protein wet-lab call. *(ii) Heterogeneous tagging convention*. The cohort spans three construct architectures (76% N/C-terminal His_6_, 33% HiBit fusion (often combined with His_6_), and 23% untagged) that the public training corpora the predictors were trained on do not span; per-construct AUCs vary by ≥ 0.2 (Table S7 caption a) and do not replicate on the public externals (§2.3). Together, these two issues mean we can use this cohort to corroborate that a deployed checkpoint is not catastrophically miscalibrated for an industrial expression pipeline (Aikium uses it for exactly this), but not to distinguish between architectures or training-pool variants. None of the 1,261 scaffolds appears in any public training corpus we tested against (0% exact-match overlap with the public training pools); all sequences are processed through a tag-aware normalisation pipeline before scoring.

The released checkpoint reaches AUC 0.865 ± 0.012 on this set (three-seed mean under the protocol-aware aggregation of Table S7), corroborating competitiveness on this descriptive cohort within its label-construction noise floor (§S6). Per-construct AUCs are 0.948 on His_6_-only (*N* = 542), 0.676 on HiBit (*N* = 424), and 0.991 on untagged (*N* = 295). Per-stringency three-seed-mean AUC ranges from 0.852 to 0.873 across the five centrifugation conditions, substantially tighter than the 0.10–0.15 spread of single-stringency specialists trained on the same data. NetSolP per-construct values are scored under tag-aware sequence normalisation (Methods §4.6); raw-input scoring of the same NetSolP checkpoint inflates per-construct NetSolP AUC by +0.01 to +0.02 through tag-feature memorisation.

#### The released checkpoints’ training pools are clean of this cohort

A source-pool decomposition of the canonical-147K and research-tier 229K training pools (Methods §4.1) finds zero scaffold-cohort proteins in either; the cohort is held out only, never used in training or validation by any released checkpoint.

---

**Table S5:**
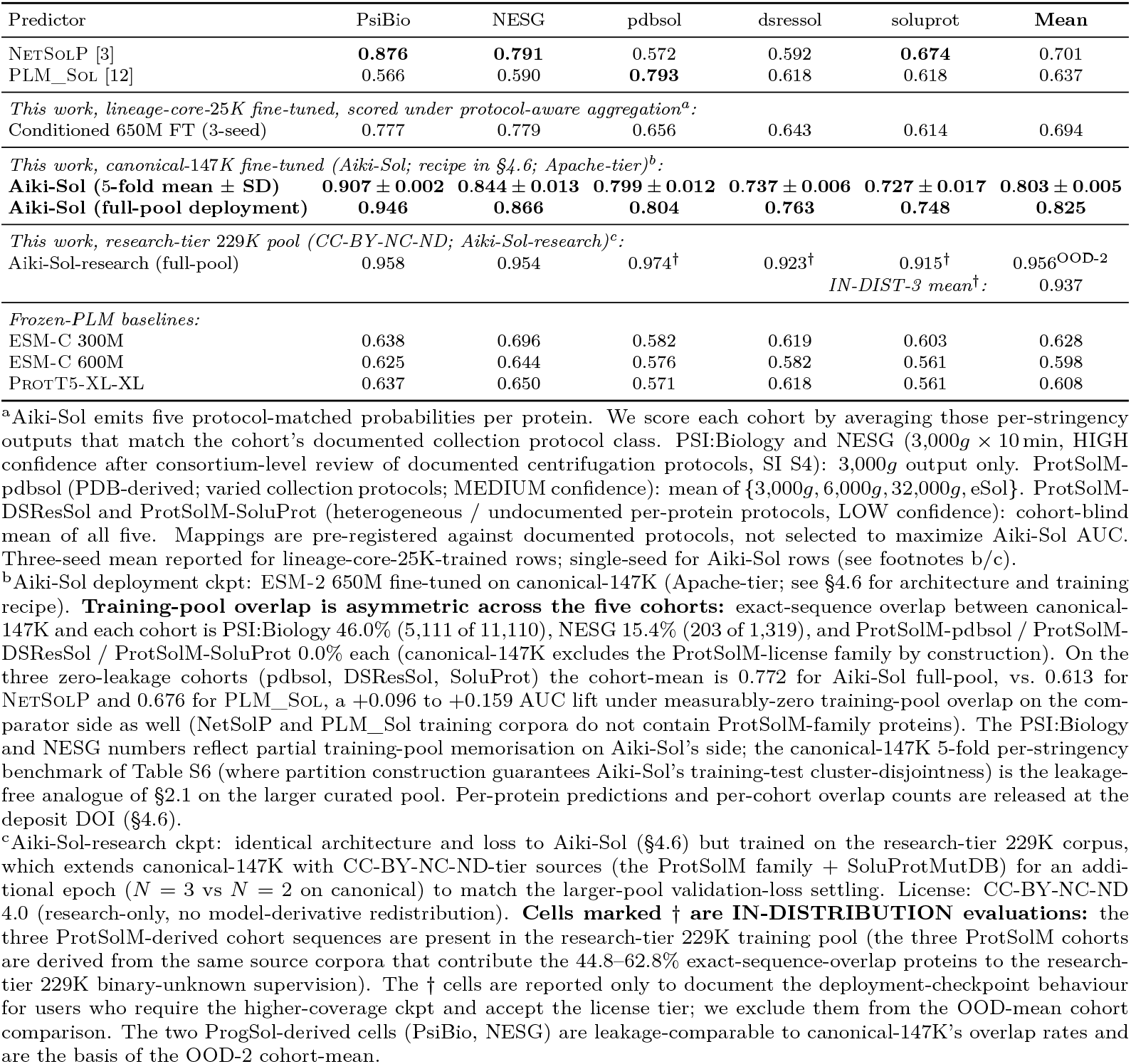
Cluster-disjoint external benchmarks at 25% identity. Aiki-Sol variants are scored under the protocol-aware aggregation of §2.3a; comparators emit a single *P* (soluble) per protein. Bold: best per column among ckpts evaluated under the same training-pool tier. The three lineage-core-25K-trained Aiki-Sol rows track within ±0.011 AUC of each other on the cohortmean (0.694–0.705) and span NetSolP’s cohort-mean of 0.701, with the ranking inversions across cohorts tracking training-distribution match more closely than any architectural advantage. The canonical-147K-trained Aiki-Sol deployment row lifts the cohort-mean by +0.12 AUC over the lineage-core-25K rows and the binary comparators on every cohort, with the largest gain on the two ProtSolM-derived cohorts whose training-pool match shifts most under the larger curated pool. The 5-fold mean ± SD row (cohort-mean 0.803 ± 0.005) shows consistent across-fold behaviour. The CC-BY-NC-ND-tier Aiki-Sol-research (research-tier 229K full-pool) row reaches OOD-mean AUC 0.956 (mean of the two training-pool-disjoint cohorts: PSI:Biology and NESG); its three ProtSolM-derived cohort cells (marked *†*) sit at AUC 0.915– 0.974 but reflect training-pool memorisation rather than generalisation (research-tier 229K includes the ProtSolM-license family during training) and are reported only to document the deployment-checkpoint’s in-distribution behaviour; we exclude them from the cohort-mean comparison against the Apache-tier deployment row. The architecture and training recipe of Aiki-Sol and Aiki-Sol-research are described in §4.6.

**Table S6:** Canonical-147K cluster-disjoint per-stringency AUC, 5-fold mean ± sample SD. Aiki-Sol architecture and training recipe are described in §4.6; comparator rows re-bin existing per-protein predictions from the protocol-stratified-85K benchmark of §2.1 under the canonical-147K fold assignment (100% coverage for 4*/*5 tools; 99.99% for SoluProt). The eSol (21,600*g ×* 30 min PURE) stringency is excluded from this table because canonical-147K routes eSol proteins to a continuous-regression target rather than a binary label; per-stringency AUC is reported on the four binary stringencies. “Mean of means” is the simple average across the four stringencies.

| Predictor | 3,000g | 6,000g | 32,000g | 100,000g | Mean of means |
| --- | --- | --- | --- | --- | --- |
| **Aiki-Sol (canonical-147K, 5/5)** | **0.762 ± 0.017** | **0.798 ± 0.017** | **0.647 ± 0.041** | **0.621 ± 0.004** | **0.707 ± 0.015** |
| NETSOLP [3] | 0.751 ± 0.011 | **0.450 ± 0.029** | 0.550 ± 0.034 | 0.525 ± 0.009 | 0.569 ± 0.128 |
| PLM_SOL [12] | 0.679 ± 0.011 | 0.756 ± 0.006 | 0.545 ± 0.032 | 0.580 ± 0.015 | 0.640 ± 0.096 |
| SoluProt [11]<sup>a</sup> | 0.653 ± 0.009 | 0.705 ± 0.014 | 0.567 ± 0.037 | 0.552 ± 0.009 | 0.619 ± 0.073 |
| DSResSol [17] | 0.615 ± 0.017 | 0.711 ± 0.017 | 0.585 ± 0.040 | 0.561 ± 0.014 | 0.618 ± 0.066 |
| DeepSol1 [8] | 0.539 ± 0.020 | 0.578 ± 0.014 | 0.566 ± 0.046 | 0.524 ± 0.009 | 0.552 ± 0.025 |

<sup>a</sup>SoluProt scored with mmseqs2 substituted for USEARCH; 3 rows have NaN predictions due to alignment failures on highly-divergent sequences (per the 0.087% failure rate documented in §2.1 footnote a).
NETSOLP’s 6,000g AUC of 0.450 ± 0.029 is below chance on all 5 folds under canonical-147K binning. This is a different stringency than the 32,000g-below-chance pattern documented in §2.1 on the protocol-stratified 85K benchmark. The change in which stringency falls below chance under different fold compositions is itself evidence that a single binary predictor cannot maintain calibration across stringencies, and that the failure is sensitive to the specific protein composition of each stringency stratum. This is the failure mode the per-protocol architecture of Aiki-Sol is designed to address. eSol-stratum AUCs are not in this table (canonical-147K eSol proteins carry continuous `continuous_eSol` labels routed to a regression head; Aiki-Sol Spearman $\rho$ = 0.574 ± 0.024 on those proteins).

**Table S7:**
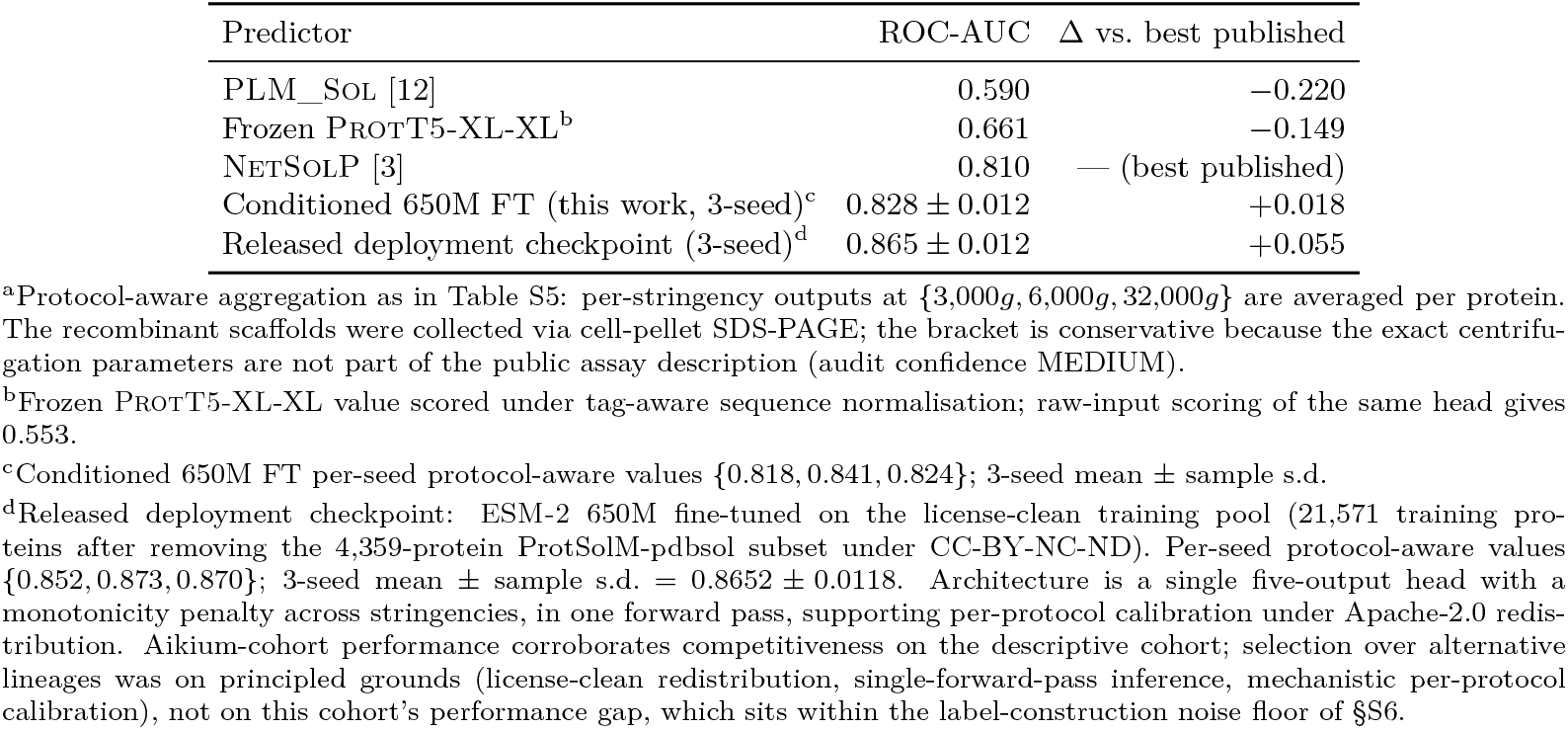
Prospective recombinant scaffold deployment (*N* = 1,261, inclusion-body versus cytosolic). All predictors scored on the same tag-normalised sequences under the protocol-aware aggregation^a^ (recombinant *E. coli* BL21(DE3) cell-pellet SDS-PAGE; protocol bracket 3,000*g*+6,000*g*+32,000*g*; MEDIUM confidence).

## References

[1] Braun, P. & LaBaer, J. High throughput protein production for functional proteomics. Trends in Biotechnology 21, 383–388 (2003).

[2] Lin, Z. et al. Evolutionary-scale prediction of atomic-level protein structure with a language model. Science 379, 1123–1130 (2023).

[3] Thumuluri, V. et al. NetSolP: predicting protein solubility in E. coli using language models. Bioinformatics 38, 941–946 (2022).

[4] Niwa, T. et al. Bimodal protein solubility distribution revealed by an aggregation analysis of the entire ensemble of Escherichia coli proteins. PNAS 106, 4201–4206 (2009).

[5] Price, W. N. et al. Understanding the physical properties that control protein crystallization by analysis of large-scale experimental data. Nature Biotechnology 27, 51–57 (2009).

[6] Acton, T. B. et al. Preparation of protein samples for NMR structure, function, and small-molecule screening studies. Methods Enzymol. 493, 21–60 (2011).

[7] Joachimiak, A. High-throughput crystallography for structural genomics. Curr. Opin. Struct. Biol. 17, 359–366 (2007).

[8] Khurana, S. et al. DeepSol: a deep learning framework for sequence-based protein solubility prediction. Bioinformatics 34, 2605–2613 (2018).

[9] Howard, J. & Ruder, S. Universal language model fine-tuning for text classification. Proc. ACL, 328–339 (2018).

[10] Steinegger, M. & Söding, J. MMseqs2 enables sensitive protein sequence searching for the analysis of massive data sets. Nature Biotechnology 35, 1026–1028 (2017).

[11] Hon, J. et al. SoluProt: prediction of soluble protein expression in Escherichia coli. Bioinformatics 37, 23–28 (2021).

[12] Zhang, X., Hu, X., Zhang, T., Yang, L., Liu, C., Xu, N., Wang, H. & Sun, W. PLM_Sol: predicting protein solubility by benchmarking multiple protein language models with the updated Escherichia coli protein solubility dataset. Briefings in Bioinformatics 25, bbae404 (2024).

[13] Li, B. & Ming, D. GATSol, an enhanced predictor of protein solubility through the synergy of 3D structure graph and large language modeling. BMC Bioinformatics 25, 204 (2024).

[14] Tan, Y., Zheng, J., Hong, L. & Zhou, B. ProtSolM: protein solubility prediction with multimodal features. Proceedings of the 2024 IEEE International Conference on Bioinformatics and Biomedicine (BIBM) 223–232 (2024).

[15] Wang, C. & Zou, Q. Prediction of protein solubility based on sequence physicochemical patterns and distributed representation information with DeepSoluE. BMC Biology 21, 12 (2023).

[16] Chen, J., Zheng, S., Zhao, H. & Yang, Y. Structure-aware protein solubility prediction from sequence through graph convolutional network and predicted contact map. Journal of Cheminformatics 13, 7 (2021).

[17] Madani, M., Lin, K. & Tarakanova, A. DSResSol: a sequence-based solubility predictor created with dilated squeeze excitation residual networks. Int. J. Mol. Sci. 22, 13555 (2021).

[18] Schmirler, R., Heinzinger, M. & Rost, B. Fine-tuning protein language models boosts predictions across diverse tasks. Nature Communications 15, 7407 (2024).

[19] Pimtawong, T., Ren, J., Lee, J.Lee, H.-M. & Na, D. A review on computational models for predicting protein solubility. Journal of Microbiology 63, e2408001 (2025).

[20] Baranowski, C., Garcia Martin, H., Oyarzún, D. A. et al. Can protein expression be “solved”? Trends in Biotechnology 43, 2724–2742 (2025).

[21] Xu, J., Wu, T., Jiang, Y., Nie, L. & Lyu, Q. MMSol: predicting protein solubility with an antinoise multimodal deep model. Journal of Chemical Information and Modeling 65, 7238–7251 (2025).

[22] Qian, J., Yang, L., Wang, R. & Qi, Y. Pro4S: prediction of protein solubility by fusing sequence, structure, and surface. bioRxiv 2025.11.05.686869 (2025).

[23] Sormanni, P., Aprile, F. A. & Vendruscolo, M. The CamSol method of rational design of protein mutants with enhanced solubility. J. Mol. Biol. 427, 478–490 (2015).

[24] Garcia-Fruitós, E., Martínez-Alonso, M., González-Montalbán, N., Valli, M., Mattanovich, D. & Villaverde, A. Divergent genetic control of protein solubility and conformational quality in Escherichia coli. J. Mol. Biol. 374, 195–205 (2007).

[25] Villaverde, A. & Carrió, M. M. Protein aggregation in recombinant bacteria: biological role of inclusion bodies. Biotechnology Letters 25, 1385–1395 (2003).

[26] EvolutionaryScale Team. ESM Cambrian: revealing the mysteries of proteins with unsupervised learning. EvolutionaryScale model release, 2024. https://www.evolutionaryscale.ai/blog/esm-cambrian and https://github.com/evolutionaryscale/esm.

